# Modularity, ecology, and theoretical evolution of the ribozyme body plan

**DOI:** 10.64898/2026.03.07.710320

**Authors:** Ido Bachelet

**Affiliations:** The Scojen Institute for Synthetic Biology, Reichman University, Herzliya, Israel

## Abstract

Ribozymes are relics of molecular life forms from the primitive earth that are embedded within modern genomes across all kingdoms of life. Despite significant knowledge from decades of bioinformatic and biochemical research, a gap remains in our understanding of the world in which ribozymes existed, their interactions, ecology, and possibly also evolution. The present study proposes a new theoretical basis for understanding these aspects of ribozyme biology by adopting a zoological frame of thought. Seven families of small self-cleaving ribozymes are each mapped to a primitive marine animal analog based on topological architecture, and classified into body plan grades paralleling cnidarian, ctenophore, and bilaterian organization. A formal notation describing ribozyme regions as bodies, cavities, and limbs enables systematic comparison with animal body plans and highlights reusability of parts across ribozyme groups, in turn enabling the construction of a connectivity network and a putative body plan-based evolutionary ordering. This ordering of body plans identifies systematic gaps corresponding to undiscovered ribozyme forms, one of which, a planktonic form of hammerhead, was bioinformatically found in 16.2% of all hammerhead sequences. Computational cross-cleavage analysis across all 49 pairwise interactions (including conspecific) suggests that the hammerhead was a generalist apex predator in the RNA world, while the hatchet was a vulnerable, filter-feeding or scavenger prey species. Conspecific analysis suggests that cannibalism was also a prevalent feeding strategy. Evolutionary avoidance signatures suggest ancient predator–prey coevolution. This theory emphasizes behavior, modularity, and ecological interactions as primary drivers of early ribozyme evolution, offering a new pathway for inferring ancient RNA forms independent of sequence-first assumptions.

## Introduction

The discovery of catalytic RNA, or ribozyme ^1,2^, was a watershed event that shifted the traditional view on RNA as a messenger, mediator, and scaffold (mRNA, tRNA, rRNA) within the central dogma of molecular biology ^3^. It confirmed the hypotheses of early thought pioneers like Rich ^4^, Woese ^5^, Crick ^6^, and Orgel ^7^, and formed the solid basis for the RNA world hypothesis ^8,9^ which is accepted as a leading theory on the origin of life. Together with other lines of evidence, including many basic molecules of biochemistry (ATP, NAD, CoA) being essentially RNA derivatives, and RNA viruses that are independent of DNA, the ribozyme highlights a hypothetical period of likely tens to hundreds of millions of years ^10,11^ during which RNA initiated the chemical reactions that would shape biology and particularly the genetic code, and underwent the first steps of evolution by natural selection.

Ribozymes have been studied extensively in the past decades, with nearly 15 groups of naturally occurring ribozymes identified to date, and *in-vitro* evolution experiments producing approximately 50 specific artificial ribozymes and likely hundreds of catalytic cores. Today naturally occurring ribozymes are found in genomes across all kingdoms of life, often embedded within larger RNA transcripts and can self-cleave or catalyze RNA processing reactions, in some cases contributing to the regulation of gene expression ^12^. Much knowledge about the structure, kinetics, and bioinformatics of ribozymes has been achieved to date. However, the period during which ribozymes dominated the world as autonomous organisms, and how they existed, behaved, and reproduced, remain mysterious.

Animals appeared on earth between 500 and 600 million years ago, and it is estimated that approximately 99% or more of the species that evolved since then have become extinct ^13^. Nevertheless, the world during the periods these animals lived in is understood to us, and much about the behavior, ecology, and evolution of extinct species has been inferred from the fossil record ^14^, particularly from animal body plans. In zoology and evolutionary biology, the concept of a body plan (*Bauplan*) refers to a recurring organizational blueprint that defines how an organism is constructed, how it interacts with its environment, and what behaviors it can express. Body plans are defined not by molecular composition but by higher-order architectural features such as symmetry, the arrangement of functional appendages, etc. This abstraction has proven essential for understanding locomotion, feeding strategy, habitat preference, ecological niche, functional and developmental constraints, lifestyle, and possibly also evolutionary history and relationships in extinct animal species.

In the context of ribozymes, the zoological frame of thought could be more than a mere metaphor. Ribozymes are first and foremost RNA molecules, with nucleotide sequence, folding thermodynamics, and kinetics of catalysis; but this view might overlook the existence of ribozymes as organisms in a prebiotic ecological system, in which they behaved, interacted, and reproduced much like animals do today. Many attempts to create self-reproducing RNA molecules have been reported to date; however, these arguably were not modeled after an animal but, rather, a polymerase.

This study extends the concept of body plans to ribozymes by viewing them as minimal animals with functional bodies whose behaviors emerge from their organization. This perspective is motivated by several lines of empirical evidence. For example, modern ribozymes, which are typically locked as single chains in genomes and operate in *cis*, can be assembled successfully from short parts ^15^, are active as assembled complexes ^16^, and are capable of operating in *trans* ^17,18^, supporting their hypothetical ancient autonomy and modular organization. Recent theoretical and experimental work increasingly supports the view that early life did not arise from a single, highly optimized RNA molecule, but rather from interacting networks of simpler components. Finally, it has recently been shown that ribozymes can form a rudimentary food chain and reuse parts from other ribozymes for reproduction ^19^.

The present study first establishes a metaphorical mapping between ribozymes and primitive marine animals, grounded in a formal notation that describes ribozyme regions as bodies (paired stems), limbs (unpaired strands), and cavities (catalytic cores). This notation enables the classification of seven families of small self-cleaving ribozymes into a set of structural body plans, and allows direct comparison with animal body plans described in the same terms. The body plans suggest reusability of parts across ribozyme groups, enabling construction of a connectivity network and a putative body plan-based evolutionary ordering. This inferred ordering is compared to established evolutionary relationships among primitive marine animals, revealing unexpected similarity across these scales. Bioinformatic validation of one predicted species, a planktonic “medusa”-like hammerhead, and computational cross-cleavage analysis across all pairwise ribozyme interactions, are presented as evidence that the body plan framework yields testable and productive hypotheses about the RNA world.

## Results

As a first step in adopting a zoological frame of thought for a ribozyme body plan-based theory, ribozymes were artistically rendered as organic, biological-like forms slightly reminiscent of Desmond Morris’ biomorphs **(Figure 1)**. Ribozymes were seriocomically assigned binomial nomenclatures, with the genus being termed *Catalyrna* (catalytic RNA). These 3D-like renders are based on predicted secondary RNA folds and not on the solved 3D structures of these ribozymes. Structural studies of ribozymes usually aim to capture active, substrate-bound states, which are indeed complex (e.g. pseudoknots, kissing loops, etc) but from a zoological perspective do not define the organisms, only a position they assume when eating.

**Figure 1.**
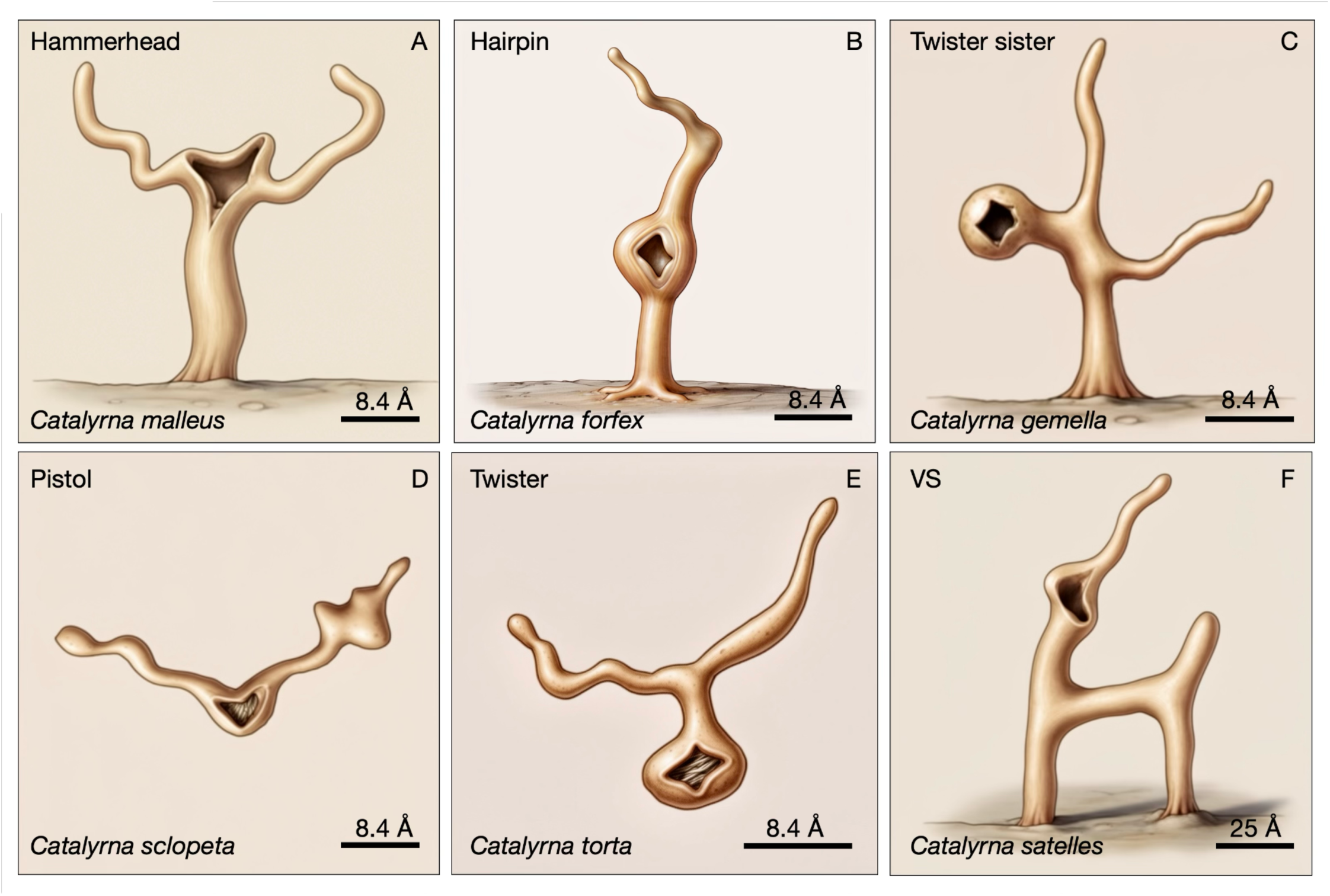
Artistic renderings of representative small self-cleaving ribozymes. Structures are shown in their native, substrate-free, *trans*-acting forms to emphasize intrinsic architectural organization rather than the catalytic conformation adopted during cleavage. Each ribozyme is represented as a biomorph-like organism and assigned a binomial-style nomenclature within the fictional genus *Catalyrna*. Panels: (A) hammerhead (*Catalyrna malleus*), (B) hairpin (*C. forfex*), (C) twister sister (*C. gemella*), (D) pistol (*C. sclopeta*), (E) twister (*C. torta*), and (F) VS ribozyme (*C. satellites*). The visual forms reflect the topology of stems, loops, and catalytic regions interpreted under the Body–Limb–Cavity framework. Scale bars indicate approximate structural dimensions derived from RNA secondary structures. These representations reflect the zoological framework used throughout the study to interpret ribozymes as architectural body plans with ecological behaviors.

Next, to establish a formal mapping between ribozymes and animals, we defined a notation that describes both in the same terms. Every RNA region falls into one of three functional types: Body (B): internally paired stems that constrain geometry, the structural scaffold; Cavity (C): the conserved unpaired regions forming the catalytic core, analogous to a mouth; and Limb (L): unpaired, non-catalytic regions that extend outward to interact with substrate, analogous to tentacles or appendages that interact with the substrate, which is analogous to food or prey. This notation deliberately suppresses sequence detail while preserving behavioral and architectural meaning.

Each of seven known self-cleaving ribozyme groups was then assigned to a primitive marine animal analog based on three criteria: (1) topological arrangement of Body, Limb, and Cavity elements in the free, *trans*-acting form; (2) positional analogy of the catalytic site to the animal’s cavity (mouth/anus); and (3) ecological lifestyle implied by the architecture (sessile vs. planktonic, ambush vs. active predator). It is hypothesized here that ribozymes with a long (⪭6 bp) body were sessile, attached to mineral surfaces via the body’s blunt end, when conditions favored and stabilized this interaction.

1. Hammerhead: *Hydra vulgaris* (solitary hydrozoan polyp). The hammerhead is a Y-shaped junction with the catalytic core (CUGAUGA…GAAA) at the tip. In the *trans* form, the catalytic cavity sits at the apex surrounded by unpaired limb strands radiating outward, identical to the hydra’s topology: a sessile polyp with tentacles and mouth at the tip. The hammerhead’s NUH↓ cleavage specificity is the broadest of any self-cleaving ribozyme, covering 12 of 16 possible dinucleotide combinations, matching the Hydra’s generalist diet.
2. Hairpin & Hatchet: *Cerianthus membranaceus* (tube anemone). The catalytic site of the hairpin and hatchet ribozymes is positioned mid-body, with paired stems extending both below and above it, and an unpaired limb beyond the cavity. Tube anemones live in tubes embedded in sediment, with the mouth positioned mid-body and tentacles extending beyond. The hairpin’s cleavage specificity (*GUC↓) is relatively narrow, consistent with a selective feeder/grazer. The hatchet’s restricted cleavage specificity (CU↓) and high fraction of unpaired residues (62%) suggest it was, too, a filter feeder that made an excellent prey.
3. Twister: *Mnemiopsis leidyi* (lobate ctenophore). Body with a central cavity and multiple arms radiating outward, closely resembling a small hydromedusa — the free-swimming medusa stage of a hydrozoan cnidarian. *Sarsia* has a prominent manubrium (feeding organ) and radiating tentacles, matching the twister’s central catalytic site and outward-projecting stems. The twister’s UA↓ cleavage specificity makes it a planktonic active predator.
4. Twister sister: *Obelia geniculata* (colonial hydrozoan). A sessile body attached to substrate by a thick stalk, with a central cavity (mouth) and multiple arms radiating from the head. Maps to Tubularia, a solitary hydroid polyp with a whorl of tentacles radiating from the body atop a stalk — the same topology. Twister sister shares UA↓ cleavage with twister but adds a basal attachment stem that anchors it, i.e. sessile rather than planktonic. Together, twister and twister sister recapitulate the cnidarian polyp–medusa duality: the same body plan in sessile and planktonic forms.
5. Pistol: *Mnemiopsis leidyi* (warted comb jelly). A central planktonic body with the catalytic core at its center and two asymmetric feeding appendages. The pistol’s GU↓ cleavage specificity is moderately broad, suggesting a planktonic ambush hunter.
6. VS ribozyme: *Haliclystus auricula* (stalked jellyfish). The most architecturally complex ribozyme among the small ones, is sessile and attached to the surface by two stems, extending from a central body. *Haliclystus* is a stauromedusa: sessile, attached by a peduncle, with multiple arms radiating from an architecturally complex body. The VS ribozyme’s G↓A/U cleavage is dependent and narrow, suggesting a sit-and-wait predator with high selectivity, not a passive filter feeder processing whatever flows by.

The seven ribozymes fall into body plan categories that parallel the major grades of primitive marine animal organization **(Figure 2)**. Diversity in hybridization lengths and melting temperatures led to the selection of specific species of ribozymes fit for specific environmental niches.

- Grade I: Cnidarian-grade body plans. Radial or near-radial symmetry, blind-sac cavity (single opening). This grade includes six of the seven known ribozymes: Plan A (solitary polyp / tip-mouth): hammerhead → *Hydra*. Plan B (anthozoan / mid-body mouth): hairpin and hatchet, a sister-species pair sharing the same four-stem, mid-body catalytic site topology but differing in exposure and ecological vulnerability. Plan C (solitary hydroid polyp): twister sister → *Tubularia*. Plan D (hydromedusa / planktonic): twister → *Sarsia*. Together, twister sister and twister recapitulate the cnidarian polyp–medusa duality, a second independent instance alongside the hammerhead polyp and its predicted medusa form. Plan E (stauromedusa / multi-armed): VS → *Haliclystus*.
- Grade II — Ctenophore-grade body plans (predicted). Biradial symmetry, planktonic. The pistol ribozyme is the only ribozyme we associated with this body plan.
- Grade III - Bilaterian-grade body plan (predicted). Bilateral symmetry, defined anterior–posterior axis, directional body organization. No known ribozyme currently occupies this grade. The bilaterian node on the animal phylogeny represents a predicted slot.

**Figure 2.**
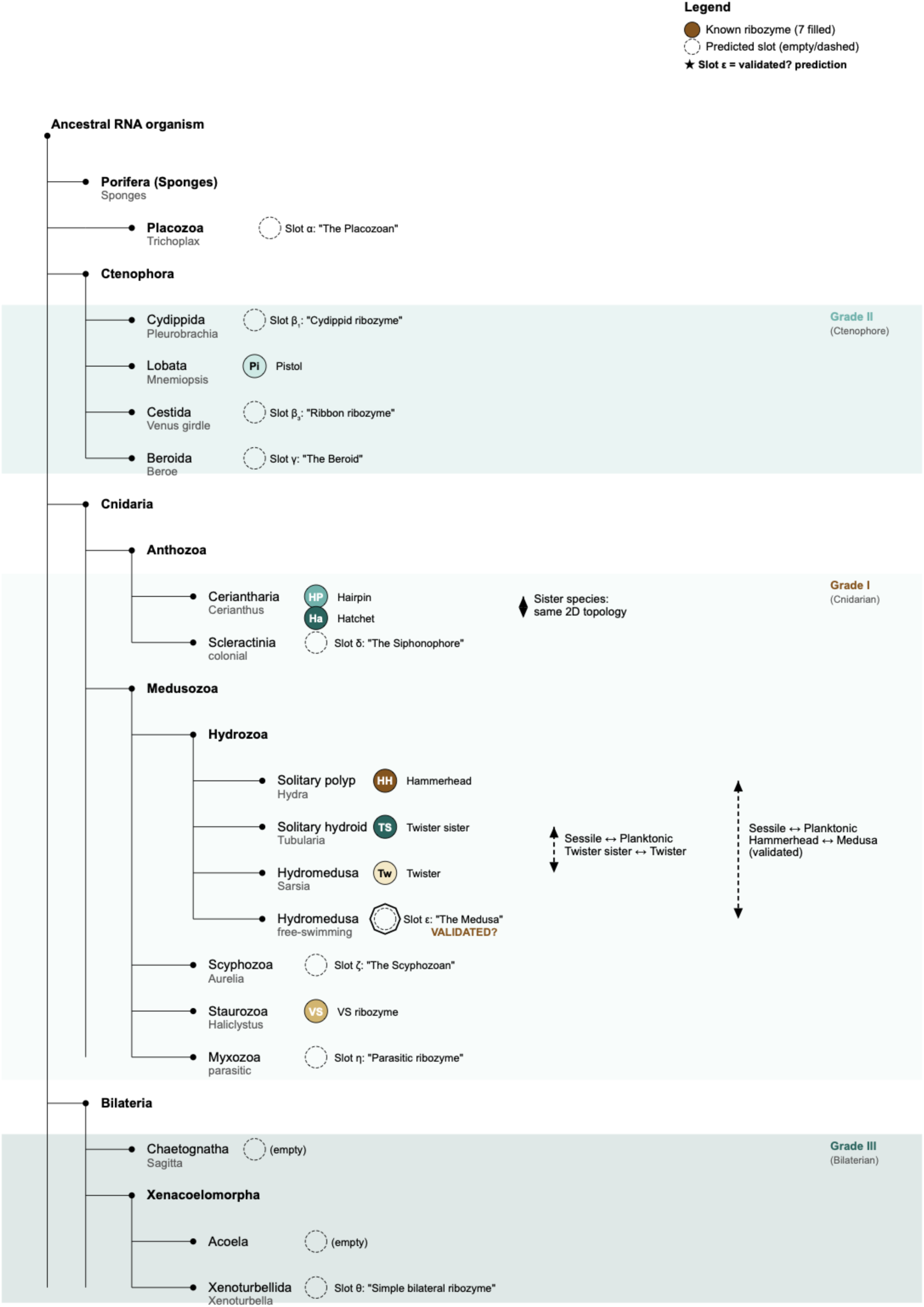
Proposed correspondence between ribozyme body plans and the major organizational grades of primitive marine animals. The diagram aligns ribozyme families with analogous metazoan body plans and places them within a conceptual evolutionary framework. Grade I (cnidarian-grade) includes most known ribozymes, characterized by radial organization and a single catalytic cavity, encompassing hammerhead, hairpin, hatchet, twister sister, twister, and VS ribozymes. Grade II (ctenophore-grade) is represented by the pistol ribozyme, displaying a biradial planktonic architecture. Grade III (bilaterian-grade) represents a predicted but currently unoccupied ribozyme body plan with bilateral symmetry and directional organization. Several nodes correspond to predicted ribozyme architectures inferred from gaps in the body-plan landscape. Arrows highlight ecological transitions between sessile and planktonic morphologies, including the predicted planktonic “medusa” form of the hammerhead ribozyme.

Weiss et al. demonstrated experimentally that a hammerhead ribozyme can function as a predator, cleaving a hairpin ribozyme’s scaffold to release a shared strand and redirect it toward hammerhead self-assembly. This proof-of-principle establishes that ribozyme food chains are biochemically feasible. Here this is extended to all seven ribozyme groups using computational cross-cleavage analysis.

To produce a qualitative description of ribozyme interactions, a dataset of 206,263 ribozyme sequences was curated **(Table S1)**. The target specificity of each ribozyme was encoded as a regular expression (hammerhead: NUH| (12 triplets); hairpin: N|GUC; twister: U|A; twister sister: U|A; pistol: G|U; hatchet: C|W; VS: G|A) and searched across all other ribozyme sequences **(Fig. 3A)**. The average number of sites per prey sequence was used as the pairwise interaction score. Based on this calculation, the hammerhead has the highest net predation score (+7.5), confirming its status as apex generalist predator. Its NUH↓ cleavage specificity, the broadest of any self-cleaving ribozyme, finds targets in every other ribozyme: 8 accessible sites in hatchet (score 16.0), 10 in VS (score 10.0), 5 in twister (score 6.5). A pairwise matrix of food chain feasibility scores was generated with these numbers **(Fig. 3B)**. Interestingly, the matrix shows that ribozymes can break down ribozymes of the same type, suggesting that cannibalism was a potential feeding strategy in the RNA world. The numbers reflect the directionality of the interaction. A corresponding food web map was plotted, in which each ribozyme is assigned a net weight (total outgoing minus total incoming) **(Fig. 3C)**. This food web is supported by computational enumeration of shared subsequences (k-mers of length ≥5 nt) across all pairwise ribozyme combinations, showing significant counts of shared k-mers. For example, hammerhead and hairpin share a 7-mer CGAAACA containing the GAAA motif - the ancestral strand demonstrated by Weiss et al. These shared parts could represent contested resources in food chain dynamics and evidence of common ancestry at the level of modular parts rather than full-length sequences.

**Figure 3.**
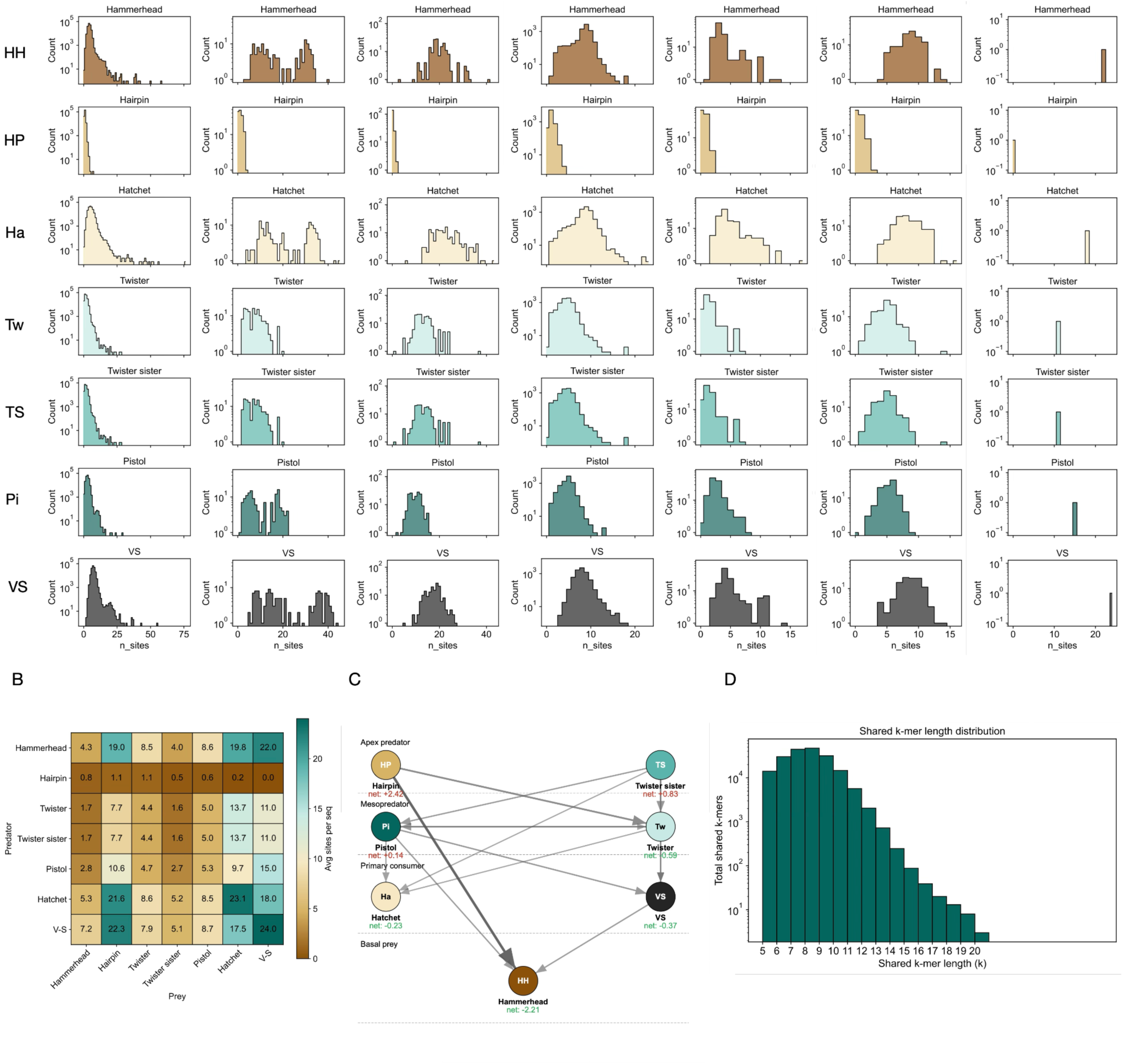
Predicted ecological interactions among ribozyme families derived from computational cross-cleavage analysis. (A) Distributions of accessible cleavage sites detected when each ribozyme motif is scanned across sequences of all other ribozymes. (B) Pairwise interaction matrix summarizing average cleavage-site counts per prey sequence, representing the feasibility of predator–prey relationships. (C) Network representation of the resulting RNA food web, where nodes represent ribozyme families and directed edges represent potential predatory cleavage interactions weighted by interaction score. Net predation values identify ecological roles within the network. The hammerhead ribozyme shows the highest positive predation score, consistent with an apex generalist predator capable of cleaving all other ribozymes. (D) Distribution of shared sequence fragments (k-mers ≥5 nt) across ribozyme families, illustrating the presence of reusable RNA modules that may act as contested resources in the molecular ecosystem.

A prediction from the evolutionary tree - the existence of a stemless “medusa” form of the hammerhead ribozyme - was tested by analyzing complete unique hammerhead sequences. For each sequence, the gap length between the conserved catalytic motifs (CUGAUGA and GAAA) was measured. The maximum possible attachment stem length was calculated as floor((gap − 3) / 2) base pairs. Sequences were classified as short-stem (gap ≤6 nucleotides, corresponding to ≤3 bp), intermediate (gap 7–10), or long-stem (gap ≥11). RNAfold was used to independently predict stem formation. The gap length distribution is strikingly bimodal **(Fig. 4)**. Of 101,757 sequences analyzed: 2,238 (2.2%) have short stems with gap ≤6 nucleotides; 434 (0.4%) fall in the intermediate range; and 99,085 (97.4%) have long stable stems with gap ≥11. The key observation is bimodality itself: the near-absence of intermediate forms (0.4%) indicates that these are two distinct populations, not a continuum. The hammerhead ribozyme exists in two discrete architectural states — a long-stemmed form and a short-stemmed form — with probable selection against intermediate configurations.

**Figure 4.**
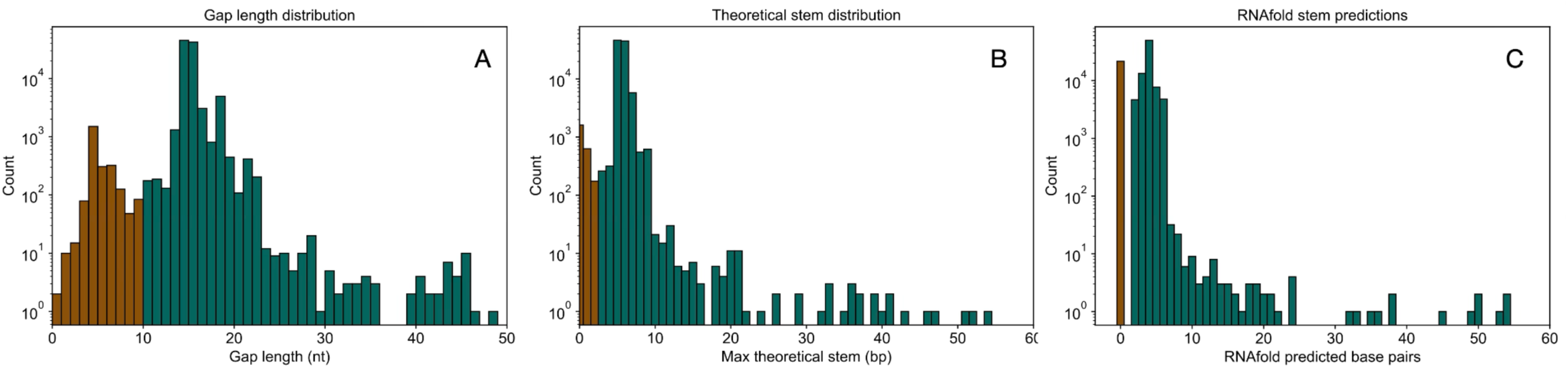
Bioinformatic analysis of hammerhead ribozyme sequences revealing two discrete architectural morphotypes. Gap lengths between conserved catalytic motifs (CUGAUGA and GAAA) were measured across 101,757 unique sequences to estimate the maximum possible attachment stem length. Sequences cluster into two dominant populations: long-stem forms (gap ≥11 nt), corresponding to a stable sessile “polyp” architecture, and short-stem forms (gap ≤6 nt), predicted to lack a substantial attachment stem and therefore represent a planktonic “medusa” morphotype. Intermediate configurations (gap 7–10 nt) are extremely rare, suggesting strong selection against unstable intermediate body plans. RNAfold predictions confirm that long-gap sequences typically form stable stems whereas short-gap sequences remain largely unpaired. The bimodal distribution supports the hypothesis that hammerhead ribozymes exist in two distinct structural states rather than a continuous morphological spectrum.

## Discussion

In the 1950s–1970s, molecular biology became organized around what later came to be known as the central dogma, articulated most clearly by Crick ^3^. Within the hierarchy implied by this framework, RNA was viewed primarily as an intermediary: a messenger (mRNA), mediator (tRNA), or structural scaffold for enzymatic complexes (rRNA) linking the informational stability of DNA with the catalytic versatility of proteins. The discovery of ribozymes challenged this hierarchy and provided a crucial missing piece to the hypothesis that RNA predated DNA in the context known as the RNA world.

Ribozymes encompass a diverse group of catalytic RNAs that vary in size, structural complexity, and biological role. Small self-cleaving ribozymes, typically ∼40–120 nucleotides long, possess compact architectures and can function as relatively autonomous catalysts. In contrast, large ribozymes such as the ribosome, RNase P, and group I and II introns span hundreds to thousands of nucleotides and form elaborate structures that function as components of complex cellular machinery ^1,2^. It is generally accepted within RNA world research that small ribozymes are far more plausible candidates as RNA world inhabitants than large structured RNAs, whose complexity likely reflects later evolutionary elaboration within cellular systems ^12,20,21^. It is important to note that the present work focuses specifically on small self-cleaving ribozymes, for this reason.

Ribozymes have been long thought of as organisms or organism analogs ^20,21^. In principle, there is no fundamental biological behavior performed by cells and animals that catalytic RNAs could not also achieve, including metabolism, replication, interaction, sensing, communication, and adaptation. Of the various properties of ribozymes, the most studied have been structure and catalysis. One possible explanation for this is that small self-cleaving ribozymes do not appear today - or, indeed, are not known to exist today - as autonomous entities free from their genomic context. A different framing could, thus, possibly enable a wider angle of observation.

This study proposes such framing by treating them as minimal organisms defined by architectural body plans. This abstraction deliberately suppresses sequence detail in favor of higher-order organizational features - bodies, limbs, and cavities - that determine how the ribozyme interacts with its environment. Such an approach parallels the use of body plans in zoology, where organisms are classified and interpreted based on structural organization rather than biochemical composition. Under this view, ribozymes become actors within a hypothetical RNA ecology, capable of behaviors such as predation, scavenging, and resource competition.

It is critical to note that the purpose of this study is not to overfit ribozymes to animals based on external similarity. Its explicit purpose is to create a metaphor on which a new framing of a scientific problem is based.

With that said, the current study faces several challenges. The main challenge is that the specific body plans described here are most likely incompatible with contemporary chemistry. For example, a planktonic form of the hammerhead ribozyme connected by a stem of 3-4 nt would be unstable in modern biology (37 °C, ∼150 mM monovalent salt, ∼1 mM divalent cations, pH ∼7.2–7.4). Yet, the prebiotic Earth likely provided diverse and extreme geochemical environments including eutectic ice phases, high monovalent salts (2–5 M) and divalent cations (1–2 M), reduced water activity on mineral surfaces, viscous or low-dielectric solvents, and a wide pH range. These conditions may have allowed RNA molecules to exhibit exotic, peculiar structures and behaviors that are rare or impossible in contemporary biological chemistry, but they cannot be fully captured by RNA folding simulations. However, employing ancient chemistry as a possible solution raises another question, since ribozyme activity is known from contemporary chemistry. This discrepancy can be resolved only by experimental investigation into the structure and catalytic kinetics of ribozymes under simulated ancient chemistry. With that said, small ribozymes are known to be active across a broad spectrum of non-biological, prebiotically plausible environments, e.g. adsorption to montmorillonite clay (Biondi et al., 2007), with Fe²⁺ instead of Mg^2+^ and in and Dead Sea pools (e.g. 30 °C, 1M Na^+^, 2M Mg^2+^) ^19,22^, in the presence of ethanol or formamide ^23^, and in high hydrostatic pressure ^24^, demonstrating that their catalytic competence extends well beyond standard physiological buffer conditions.

The ecological interpretation proposed here is further supported by the computational interaction network among ribozyme families. The hatchet ribozyme, for example, has the most negative net predation score (−11.5) and the highest proportion of unpaired residues (62%), making it particularly vulnerable to trans-cleavage by other ribozymes. Its modern confinement largely to Crassvirales bacteriophage genomes—only ∼210 known copies—may reflect the long-term outcome of such vulnerability, suggesting that ancient ecological pressures could have restricted it to a narrow evolutionary refuge.

A defining feature of this framework is its deliberate abstraction away from nucleotide-level sequence detail. Whereas most studies of ribozyme evolution emphasize sequence conservation, mutational pathways, and thermodynamic optimization, the present approach operates primarily at the level of behavioral morphology - the spatial organization of functional regions and their interactions. In a primitive RNA world, it is plausible that function emerged initially from coarse-grained architectural arrangements, with specific nucleotide sequences evolving later to stabilize and refine these arrangements. In this view, sequence serves as a flexible substrate that adapts to preserve functional body plans rather than as the primary driver of early evolutionary innovation.

An important implication of the body-plan framework is that it generates testable predictions independent of sequence-based evolutionary reconstruction. The predicted existence of a planktonic “medusa” form of the hammerhead ribozyme represents one such example. The observed bimodal distribution of catalytic-core spacing among hammerhead sequences supports the existence of two discrete architectural states corresponding to sessile and planktonic variants. More broadly, the body-plan ordering proposed here predicts structural gaps corresponding to ribozyme architectures that have not yet been identified. Continued bioinformatic exploration of genomic and metagenomic datasets may therefore reveal additional ribozyme families occupying currently empty positions in the proposed morphological landscape.

The ecological interpretation also suggests that ribozyme evolution may have been driven not only by catalytic efficiency but by interaction dynamics within molecular communities. Predation, competition for reusable RNA fragments, and cooperative assembly of functional complexes could all contribute selective pressures shaping ribozyme architectures. Such dynamics resemble ecological processes observed in modern biological systems and may represent an early stage of Darwinian evolution in which selection acted on networks of interacting molecules rather than on isolated replicators.

Finally, the framework presented here should be regarded primarily as a conceptual model rather than a literal reconstruction of the RNA world. The zoological analogies used throughout are intended to illuminate structural and behavioral relationships among ribozymes rather than to imply direct evolutionary continuity between RNA molecules and animal organisms. Nevertheless, by reframing ribozymes as minimal organisms with body plans, ecological roles, and behavioral repertoires, the approach highlights a neglected dimension of early molecular evolution. It suggests that the emergence of life may have involved not only the optimization of catalytic molecules but also the appearance of primitive ecological systems in which molecular organisms interacted, competed, and diversified.

## Methods

### Body–Limb–Cavity (B-L-C) notation

A formal notation, termed Body–Limb–Cavity (B-L-C), was developed to describe ribozyme secondary-structure architecture using the same descriptive terms applicable to animal body plans. Three element types are defined. Body (B) denotes internally paired stems — double-stranded RNA helices that constrain the overall geometry of the ribozyme fold, analogous to the body wall. Limb (L) denotes unpaired, non-catalytic single-stranded regions that extend outward from the structural core to engage a substrate RNA molecule in trans, analogous to tentacles or appendages. Cavity (C) denotes conserved unpaired regions that constitute the catalytic core, analogous to a mouth or gastrovascular cavity. The notation deliberately suppresses nucleotide-sequence detail while preserving topological and behavioral meaning.

The B-L-C formula for each ribozyme was constructed by reading the secondary structure of the *trans-acting* strand (catalytic strand alone, no substrate bound) from 5′ to 3′ and recording each element in the order encountered; adjacent elements of the same type were collapsed into a single symbol. All ribozymes were analyzed in the substrate-free trans-acting form in order to describe the intrinsic body plan rather than the conformation adopted during catalysis.

### Ribozyme–animal assignments and body plan grades

Each of the seven known small self-cleaving ribozyme families was assigned to a primitive marine animal analog. Assignments were based on the convergence of three independent criteria applied to the trans-acting form: (1) topological arrangement of B, L, and C elements; (2) positional analogy of the catalytic site to the animal’s oral or gastrovascular opening; and (3) ecological lifestyle implied by the architecture (sessile vs. planktonic; ambush vs. active predation). Ribozymes bearing a long attachment stem (>6 bp) were treated as sessile, hypothesized to anchor to mineral surfaces via the stem’s blunt end. An assignment was accepted only when all three criteria were satisfied simultaneously.

Ribozymes were further organized into body plan grades paralleling the major grades of primitive marine animal organization. Grade I (cnidarian) encompasses families with radial or near-radial symmetry. Grade II (ctenophore) encompasses families with biradial symmetry. Grade III (bilaterian) corresponds to bilateral symmetry with a defined anterior–posterior axis. A Mendeleev-style classification table was constructed with B-L-C body plan categories as columns and grades I–III as rows. Cells lacking a known ribozyme representative were designated as predicted slots (α–θ) and assigned confidence levels (high / moderate / low) based on: the occupancy of adjacent cells in the table; the RNA structural plausibility of the predicted B-L-C formula; and the existence of animal body plans corresponding to the predicted category that currently lack a ribozyme counterpart.

### Modularity analysis and evolutionary overlay

Each ribozyme family was decomposed into its constituent B, L, and C elements. Shared subsequences (k-mers of length ≥5 nt) were enumerated across all 21 pairwise family combinations, and pairwise k-mer intersection counts were computed. These counts served as a measure of sequence-level raw material potentially available for modular exchange between families.

A ribozyme body plan tree was constructed from the resulting modularity network, with each family represented as a node and edges weighted by k-mer overlap and B-L-C module compatibility. The tree was rooted at the family with the highest modularity score and simplest B-L-C formula. This ribozyme tree was then overlaid onto the consensus metazoan phylogeny (Porifera → Placozoa → Ctenophora → Cnidaria → Bilateria) by aligning each ribozyme’s animal analog to its corresponding phylogenetic position, and the two independent orderings were compared.

### Sequence dataset

Ribozyme sequences were obtained from Rfam release 15.1 (downloaded 3 March 2026; ftp://ftp.ebi.ac.uk/pub/databases/Rfam/15.1/). Accessions used: hammerhead — RF00163, RF02276, RF00008, RF02275, RF02277, RF03152; hairpin — RF00173, RF04190, RF04191 (Weinberg *et al.*, 2021, DOI 10.1093/nar/gkab454); twister — RF03160, RF03154, RF02684; twister sister — RF02681; pistol — RF02679; hatchet — RF02678. The VS ribozyme sequence was obtained from PDB 4R4V (*Neurospora crassa* VS ribozyme, 186 nt), as this family is not well represented in Rfam. Sequences were extracted from Stockholm-format alignment files; gap characters were removed, thymine was converted to uracil, and duplicates were removed by both genomic coordinates and sequence identity.

### Cross-cleavage analysis

A computational food web was constructed by evaluating all 49 pairwise cleavage interactions (7 predator × 7 prey families, including the 7 conspecific diagonal entries). The cleavage specificity of each family was encoded as a regular expression for computational scanning: hammerhead NUH↓ — [ACGU]U[ACU]; hairpin N|GUC↓ — [ACGU]GUC; twister U|A — UA; twister sister U|A — UA; pistol G|U — GU; hatchet C|W↓ (W = A or U) — C[AU]; VS G|A — GA. Each predator motif was scanned across all prey sequences in a 5′-to-3′ overlapping window. The average number of matching sites per prey sequence was used as the pairwise interaction score, normalizing for differences in family size.

Accessibility filtering was applied to exclude cleavage sites buried within base-paired regions of the folded prey molecule. Secondary-structure prediction was performed for each prey sequence using RNAfold (ViennaRNA package; Lorenz *et al.*, 2011; 37°C, Turner 2004 energy parameters). A cleavage site was scored as accessible only if all nucleotide positions of the recognition motif fell within unpaired regions of the predicted minimum free energy (MFE) structure; sites with any base-paired motif positions were excluded from the accessible site count.

A composite food chain score was computed for each directed predator–prey interaction as:

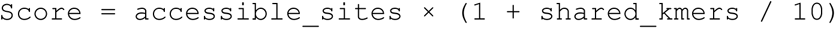

where *accessible_sites* is the number of accessible cleavage sites in the prey sequence and *shared_kmers* is the total count of k-mers of length ≥5 nt shared between predator and prey sequences. The net predation score for each family was computed as:

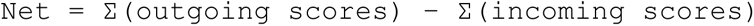

Conspecific (cannibalism) scores were computed identically, with predator and prey drawn from the same family.

An observed/expected (O/E) ratio analysis was performed to assess whether cleavage motif frequencies in prey sequences deviate from random expectation. For a motif of length *m* in a prey sequence of length *n*, the expected count under a mononucleotide null model was computed as E = n × Π(f*i*) for i = 1 to m, where f*i* is the frequency of the nucleotide at motif position i in the prey sequence. O/E ratios were computed for all 49 predator–prey pairs.

### Hammerhead medusa bioinformatic analysis

To test the prediction, derived from the body plan evolutionary tree, that a stemless planktonic “medusa” form of the hammerhead ribozyme exists, attachment stem lengths were analyzed across all unique hammerhead sequences from the six Rfam accessions listed above. For each sequence, two conserved catalytic motifs were located by pattern matching: the 5′ motif (CUGAUGA; relaxed variants CUGANGA, CUGAUGU, and CUGACGA permitted) and the 3′ motif (GAAA). Gap length was defined as the number of nucleotides between the end of the 5′ motif and the start of the 3′ motif, exclusive of the motif residues themselves; when multiple matches were possible, the shortest gap was selected. The maximum possible attachment stem length was estimated as floor((gap − 3) / 2) bp, where subtraction of 3 accounts for the minimum loop size required to reverse strand direction.

Sequences were classified into three categories: short-stem (gap ≤6 nt; predicted medusa morphotype), intermediate (gap 7–10 nt), and long-stem (gap ≥11 nt; predicted polyp morphotype). Independent structural validation was performed using RNAfold (default parameters, run in batches of 500 sequences) to confirm whether the gap region formed a helical stem or remained unpaired in the predicted MFE structure.

### Software and data availability

Secondary-structure predictions were performed with RNAfold from the ViennaRNA package (Lorenz *et al.*, Algorithms for Molecular Biology 6, 26, 2011; https://www.tbi.univie.ac.at/RNA/). All computed data tables — including the 7×7 cross-cleavage interaction matrix, food chain scores, net predation scores, O/E ratios, shared k-mer counts, and B-L-C formulas for all families — are provided as supplementary materials.

## Author contributions

I.B. designed experiments, performed experiments, analyzed data, and wrote manuscript.

---

**Table S1:**

| family | unique_sequences | avg_length_nt | rfam_accessions | other_sources | rfam_release | download_date |
| --- | --- | --- | --- | --- | --- | --- |
| Hammerhead | 199244 | 46 | RF00163,<br>RF02276,<br>RF00008,<br>RF02275,<br>RF02277,<br>RF03152 |  | 15.1 | 2026-03-03 |
| Hairpin | 139 | 165.3 | RF00173,<br>RF04190,<br>RF04191 | Weinberg et al. 2021<br>(DOI 10.1093/nar/gkab454) | 15.1 | 2026-03-03 |
| Twister | 6481 | 69.1 | RF03160,<br>RF03154,<br>RF02684 |  | 15.1 | 2026-03-03 |
| Twister sister | 129 | 81.5 | RF02681 |  | 15.1 | 2026-03-03 |
| Pistol | 103 | 68 | RF02679 |  | 15.1 | 2026-03-03 |
| Hatchet | 166 | 130.1 | RF02678 |  | 15.1 | 2026-03-03 |
| VS | 1 | 186 |  | PDB 4R4V | n/a | 2026-03-03 |

**Supplementary Table 1.** Ribozyme–animal assignments and B-L-C classification.

| Ribozyme | Latin name | Animal analog | Body plan grade | Symmetry | Cleavage specificity | Ecological role |
| --- | --- | --- | --- | --- | --- | --- |
| Hammerhead | *C. malleus* | *Hydra vulgaris* | Grade I (Cnidarian) | Radial | NUH↓ | Apex predator |
| Hairpin | *C. forfex* | *Cerianthus membranaceus* | Grade I (Cnidarian) | Radial | *GUC↓ | Primary prey |
| Twister sister | *C. gemella* | *Tubularia indivisa* | Grade I (Cnidarian) | Radial | UA↓ | Basal |
| VS ribozyme | *C. satelles* | *Haliclystus auricula* | Grade I (Cnidarian) | Radial | G↓A/U | Apex specialist |
| Twister | *C. torta* | *Sarsia tubulosa* | Grade I (Cnidarian) | Radial | UA↓ | Mesopredator |
| Pistol | *C. sclopeta* | *Sagitta elegans* | Grade III (Bilaterian) | Bilateral | GU↓ | Mesopredator |
| Hatchet | *C. securis* | Hairpin sister species | Grade I (Cnidarian) | Radial | CU↓ | Basal prey |

**Supplementary Table 2.** Complete cross-cleavage interaction matrix for all 49 pairwise ribozyme interactions including conspecific (cannibalism) scores.

| Predator | Prey | Total sites | Accessible sites | Density (/100 nt) | Shared k-mers | Longest k-mer | Food chain score |
| --- | --- | --- | --- | --- | --- | --- | --- |
| hammerhead | hammerhead | 6 | 1 | 7.14 | 0 | 0 | 1.0 |
| hammerhead | hairpin | 4 | 2 | 7.69 | 0 | 0 | 2.0 |
| hammerhead | twister | 9 | 5 | 12.86 | 3 | 5 | 6.5 |
| hammerhead | twister_sister | 3 | 0 | 3.75 | 2 | 5 | 0.0 |
| hammerhead | pistol | 7 | 2 | 10.45 | 5 | 5 | 3.0 |
| hammerhead | hatchet | 12 | 8 | 15.00 | 10 | 6 | 16.0 |
| hammerhead | VS | 22 | 10 | 11.83 | 0 | 0 | 10.0 |
| hairpin | hairpin | 0 | 0 | 0.00 | 0 | 0 | 0.0 |
| hairpin | hammerhead | 2 | 1 | 2.38 | 0 | 0 | 1.0 |
| hairpin | twister | 1 | 1 | 1.43 | 0 | 0 | 1.0 |
| hairpin | twister_sister | 1 | 0 | 1.25 | 4 | 5 | 0.0 |
| hairpin | pistol | 0 | 0 | 0.00 | 6 | 7 | 0.0 |
| hairpin | hatchet | 0 | 0 | 0.00 | 5 | 5 | 0.0 |
| hairpin | VS | 0 | 0 | 0.00 | 0 | 0 | 0.0 |
| twister | twister | 5 | 2 | 7.14 | 0 | 0 | 2.0 |
| twister | hammerhead | 3 | 3 | 3.57 | 3 | 5 | 3.9 |
| twister | hairpin | 3 | 1 | 5.77 | 0 | 0 | 1.0 |
| twister | twister_sister | 1 | 1 | 1.25 | 18 | 9 | 2.8 |
| twister | pistol | 3 | 3 | 4.48 | 0 | 0 | 3.0 |
| twister | hatchet | 9 | 8 | 11.25 | 0 | 0 | 8.0 |
| twister | VS | 11 | 4 | 5.91 | 0 | 0 | 4.0 |
| twister_sister | twister_sister | 1 | 1 | 1.25 | 0 | 0 | 1.0 |
| twister_sister | hammerhead | 3 | 3 | 3.57 | 2 | 5 | 3.6 |
| twister_sister | hairpin | 3 | 1 | 5.77 | 4 | 5 | 1.4 |
| twister_sister | twister | 5 | 2 | 7.14 | 18 | 9 | 5.6 |
| twister_sister | pistol | 3 | 3 | 4.48 | 0 | 0 | 3.0 |
| twister_sister | hatchet | 9 | 8 | 11.25 | 0 | 0 | 8.0 |
| twister_sister | VS | 11 | 4 | 5.91 | 0 | 0 | 4.0 |
| pistol | pistol | 4 | 0 | 5.97 | 0 | 0 | 0.0 |
| pistol | hammerhead | 5 | 3 | 5.95 | 5 | 5 | 4.5 |
| pistol | hairpin | 4 | 0 | 7.69 | 6 | 7 | 0.0 |
| pistol | twister | 6 | 4 | 8.57 | 0 | 0 | 4.0 |
| pistol | twister_sister | 2 | 0 | 2.50 | 0 | 0 | 0.0 |
| pistol | hatchet | 3 | 1 | 3.75 | 0 | 0 | 1.0 |
| pistol | VS | 15 | 6 | 8.06 | 0 | 0 | 6.0 |
| hatchet | hatchet | 17 | 11 | 21.25 | 0 | 0 | 11.0 |
| hatchet | hammerhead | 11 | 4 | 13.10 | 10 | 6 | 8.0 |
| hatchet | hairpin | 4 | 3 | 7.69 | 5 | 5 | 4.5 |
| hatchet | twister | 9 | 4 | 12.86 | 0 | 0 | 4.0 |
| hatchet | twister_sister | 5 | 1 | 6.25 | 0 | 0 | 1.0 |
| hatchet | pistol | 7 | 3 | 10.45 | 0 | 0 | 3.0 |
| hatchet | VS | 18 | 6 | 9.68 | 0 | 0 | 6.0 |
| VS | VS | 24 | 13 | 12.90 | 0 | 0 | 13.0 |
| VS | hammerhead | 16 | 9 | 19.05 | 0 | 0 | 9.0 |
| VS | hairpin | 7 | 3 | 13.46 | 0 | 0 | 3.0 |
| VS | twister | 9 | 5 | 12.86 | 0 | 0 | 5.0 |
| VS | twister_sister | 3 | 1 | 3.75 | 0 | 0 | 1.0 |
| VS | pistol | 10 | 4 | 14.93 | 0 | 0 | 4.0 |
| VS | hatchet | 9 | 5 | 11.25 | 0 | 0 | 5.0 |

**Supplementary Table 3.** Net predation scores and ecological roles.

| Ribozyme | Net score | Trophic role | Animal analog |
| --- | --- | --- | --- |
| Hammerhead | +7.5 | Apex predator | Hydra (sessile polyp) |
| Twister | −3 | Mesopredator | Sarsia (hydromedusa) |
| Pistol | 0 | Mesopredator | Sagitta (arrow worm) |
| VS | −3 | Apex specialist | Haliclystus (stalked) |
| Hairpin | −10 | Primary prey | Cerianthus (tube anemone) |
| Hatchet | −12 | Basal prey | Hairpin sister species |
| Twister sister | +21* | Basal (anomaly) | Tubularia (solitary hydroid) |

*\*Twister sister's high score reflects sequence-level cleavage site counts; its trans-cleavage efficiency is poorly characterized and likely overestimated.*

**Supplementary Table 4.** Abundance distribution of self-cleaving ribozyme groups.

| Ribozyme | Rfam copies | Species range | Trophic role | Host range |
| --- | --- | --- | --- | --- |
| Hammerhead | ~326,852 | ~997 spp | Apex predator | All domains + viroids |
| Twister | ~6,998 | ~175 spp | Mesopredator | Bacteria + eukaryotes |
| Hairpin | ~945 | Plant viruses | Primary prey | Satellite RNAs only |
| Pistol | ~500 | Bacteria + phage | Mesopredator | Bacteria + phage |
| Hatchet | ~210 | Phage only | Basal prey | Crassvirales only |
| Twister sister | 17 | 12 spp | Basal | Bacteria + 1 archaeon |
| VS | 1 | 1 spp | Apex specialist | Neurospora mitochondria |

**Supplementary Table 5.** Hammerhead stem classification summary.

| Classification | Count | Percentage | Gap length range (nt) | Max stem (bp) |
| --- | --- | --- | --- | --- |
| Short-stem (stemless) | 435 | 16.2% | 2–6 | $\leq 3$ |
| Intermediate | 15 | 0.6% | 7–9 | 3–5 |
| Long-stem (normal) | 2239 | 83.3% | 11–70 | $\geq 4$ |
| **Total** | **2689** | **100.0%** | — | — |

**Rfam family breakdown for short-stem population**
| Rfam family | Count | Percentage |
| --- | --- | --- |
| RF00008 (Type III) | 415/435 | 95.4% |
| Other families | 20/435 | 4.6% |

